# Incidental conformational switching in an allosteric enzyme

**DOI:** 10.64898/2026.08.04.741757

**Authors:** Paul J. Sapienza, Darex J. Vera-Rodriguez, Trevor R. Mileur, Andrew L. Lee

**Author notes:** Corresponding authors: Paul J. Sapienza, Division of Chemical Biology and Medicinal Chemistry, UNC Eshelman School of Pharmacy, University of North Carolina at Chapel Hill, 4109 Marsico Hall 125 Mason Farm Rd., Chapel Hill, NC 27599-7363, Andrew L. Lee, Division of Chemical Biology and Medicinal Chemistry, UNC Eshelman School of Pharmacy, University of North Carolina at Chapel Hill, 4109 Marsico Hall 125 Mason Farm Rd., Chapel Hill, NC 27599-7363. These authors contributed equally to this work (Paul J. Sapienza, Darex J. Vera-Rodriguez).

## Abstract

The classical understanding of allostery was initially grounded in two-state models, such as MWC and KNF, where structure and function are inextricably linked through transitions between low-(T) and high-affinity (R) states. Here, we show Yeast chorismate mutase (CM) provides a vivid example of the growing list of exceptions to the traditional T vs R two-state allosteric paradigm. While CM exhibits dynamic sampling of the R-state in the presence of the activator tryptophan (Trp), suggesting a conformational selection (CS) mechanism, we present multiple instances where conformational status and catalytic activity are decoupled. Using NMR spectroscopy and kinetic assays, we identify CM variants that reside almost exclusively in the T conformation can exhibit maximal activity, while others that predominantly occupy the R conformation are weakly active. Quantitative comparison of experimental data with a parameterized CS model reveals deviations of up to two orders of magnitude, ruling out the simplest two-state model for substrate affinity modulation in this system. We propose that the observed T-to-R switching in CM is “incidental”, a byproduct of an evolved energy landscape that allows access to the substrate-bound pose but does not mechanistically determine affinity. Our findings suggest that allosteric regulation in CM may instead be driven by local features of the ground-state ensemble, which operate independently of global T/R status. This work further highlights an emerging view that the mere observation of a pre-sampled active conformation does not sufficiently prove a two-state mechanism and further underscores the need for deeper ensemble-based perspectives in protein engineering and allostery.

## Introduction

For the first 40 years after the term ‘allostery’ was coined(1), the “second secret of life” was more or less exclusively understood by the low (T) and high (R) affinity formalism in which structure and function were inextricably linked. Homo-oligomers are the largest and most well studied class of allosteric proteins(2) for which ligand binding to the first subunit makes subsequent binding of the same type of molecule to the other subunits more or less favorable. Within this oligomeric group, two phenomenological models prevailed: MWC(3) and KNF(4) distinguished by whether the structural change from T to R occurs in a concerted manner in all subunits or occurs only after binding, respectively. A related branch of this formalism is also routinely applied to monomeric systems with binding linked conformational change, in which the transition occurs prior (conformational selection, CS), or after (induced fit, IF(5)) binding. CS occurs when the unbound protein exists primarily in a binding-incompetent conformation but can exchange into a binding-competent one, which then preferentially binds ligand: i.e. ligand binding selects the active conformation. MWC and CS selection can be considered related mechanisms as they rely on exchange between low and high affinity states without substrate. Mechanistically, MWC and CS highlight proteins’ abilities to spontaneously move in ‘purposeful’ ways and advertises how proteins often need to be dynamic to fulfill their function. Upon observation of a protein’s structure moving to a different conformation, it is natural to consider this as some kind of functionally relevant state and to ascribe meaning. This has led to important insights with identification of essential functional conformations(6-10). In most cases, these exact conformations could be observed by NMR and capturing these rare and transient states provided opportunities for NMR to ‘flex’ its ability to identify functional dynamics.

Recently, a growing number of studies are adding complexity and therefore a deeper understanding of allosteric and non-allosteric mechanisms of binding. The ensemble allosteric model moves beyond the simple two state structural picture and aspires to consider the entire ensemble of states at equilibrium(11). This type of framework is required to describe cases where the allosteric mechanism differs with different ligands(12), purely entropic allostery independent of, or in the absence of conformational change(13-16), or a case in which triggering a T state is not sufficient to create the inhibitory allosteric signal(17). Adding to this modern view is a flux-based framework for distinguishing between various binding mechanisms. It considers the rates of interconversion between high and low affinity states, the on and off rates for each, and the ligand concentration(18, 19). Integration of this information is not trivial as it involves structural, kinetic, and oftentimes systems amenable to sophisticated NMR analysis. In some cases, these studies do not arrive at ‘either-or’ mechanisms, but rather flux through both CS *and* IF, depending on the condition(20, 21). Despite growing appreciation of the expanded and varied repertoire, even within systems, we are still in an early stages of both collecting examples and using accrued knowledge to design proteins with functional dynamic properties(22). This communication is concerned with whether a two state T vs. R structure-based activation model is sufficient to explain the behavior of the model dimeric allosteric enzyme, yeast chorismate mutase (CM).We previously reported that CM exhibits dynamic sampling of a second conformer, the R state, in the absence of substrate-like ligand, consistent with CS(23). CM is a homodimer that exhibits both homotropic and heterotropic allosteric function. Its function is to catalyze the conversion of chorismate to prephenate in aromatic amino acid synthesis, and it does so with positive cooperativity in the effector-free and tyrosine-bound states(24). The enzyme is activated upon binding Trp and inhibited upon binding Tyr. From x-ray crystallographic studies, CM was shown to be able to adopt T and R structures, in accordance with binding its various ligands. We added to these structures, recently, by showing that Trp binding to CM alone (TrpCM) actually retains the T structure but leads to a small (8%) sampling of the R conformation, switching dynamically on the low millisecond timescale. R-state crystal structures have an active site configuration that is sterically and electrostatically more compatible with binding (Figure S1A). Further, this switching is absent in TyrCM, leading to a reasonable hypothesis that Trp enhances substrate binding through a CS mechanism(23) and that this enzyme abides by a simple two state T and R model.

However, in several instances we have observed a poor level of correlation between CM conformational sampling of a more “active” structure and its catalytic function or regulation of that function. Specifically, from the perspective of a tight coupling between the R-state structure and affinity modulation, we have observed discordant features. These are: (1) Binding of activator Trp leaves CM primarily in the T conformation; (2) Though TrpCM is mainly T, the activity curve reflects maximal activation with clear hyperbolic saturation behavior, and, sigmoidicity would be observed if subunit switching were concerted as in the MWC model; (3) Similarly but more extreme, and shown herein, TyrCM^T226I^ exists in the T conformation (∼100%) but is fully activated; (4) We have also identified a mirror case in which a double mutant adopts the R conformation, yet is *inactive*; and (5) Despite showing an ability to adopt T or R conformations, when considering both subunits in the dimer, CM behavior appears inconsistent with strict definitions of both the MWC and KNF models. Based on these and other observations, we are forced to consider that, at least in this specific CM where activity is regulated by allosteric effectors, significant pre-sampling of an “activated” conformation might not be determinative mechanistically. Rather, the pre-sampling may merely reflect an inability to avoid the transition the enzyme has evolved to perform upon substrate recognition based on its coordinates on the energy landscape. And in these cases, it is the features of the ground state ensemble that dominate the thermodynamics of binding. Here, we review and show the data supporting this view for CM, as well as show that in cases where some level of correlation between conformation and activity are observed, the conformational change is likely to be incidental. This raises the question of why conformational switching occurs, but more importantly what as-of-yet undetected mechanisms *are* responsible for changes in protein function (in this case, substrate binding affinity)? Our view is not to dismiss the importance of conformational change; CM must adopt the R-state for activity, but it may be only a part of a larger regulatory or substrate flux mechanism. We argue that in CM, and likely other systems, it is possible to decouple T or R conformational status from the operative regulatory scheme.

## Results

### T conformation, high activity

Figure 1 shows CM activity curves for three forms of CM. Wild-type CM with Tyr bound (TyrCM) yields a sigmoidal curve, as shown previously(24), consistent with retention of positive cooperativity. By contrast, strictly hyperbolic curves with higher activity and much tighter binding are observed for both TrpCM^T226I^ and TyrCM^T226I^. T226I is a constitutively active mutant(25), presumably by stabilizing the R conformation which was confirmed for TrpCM^T226I^ x-ray structures(26). It was previously thought that T226I-CM cannot bind Tyr(27), but we found that Tyr binds T226I readily (Figure S2), and methyl NMR spectra show unequivocally that TyrCM^T226I^ is in the T conformation, without line broadening (Figure 1A). That this form adopts the T conformation is inconsistent with the high-activity, hyperbolic profile observed (Figure 1B) that is expected from a preference for the R conformation under the two-state model. We further investigated TyrCM^T226I^ by NMR to see if there was evidence for switching to R that might help to explain the activity. Methyl ^1^H CPMG curves were flat relative to TrpCM and, while there is evidence for a high energy state resembling R, its population is less than 1% (Figure S3) or 10-fold lower compared to TrpCM(23). In summary, TyrCM^T226I^ presents a case where CM is clearly in the T conformation and does not appreciably sample R yet exhibits maximal activity.

**Figure 1.**
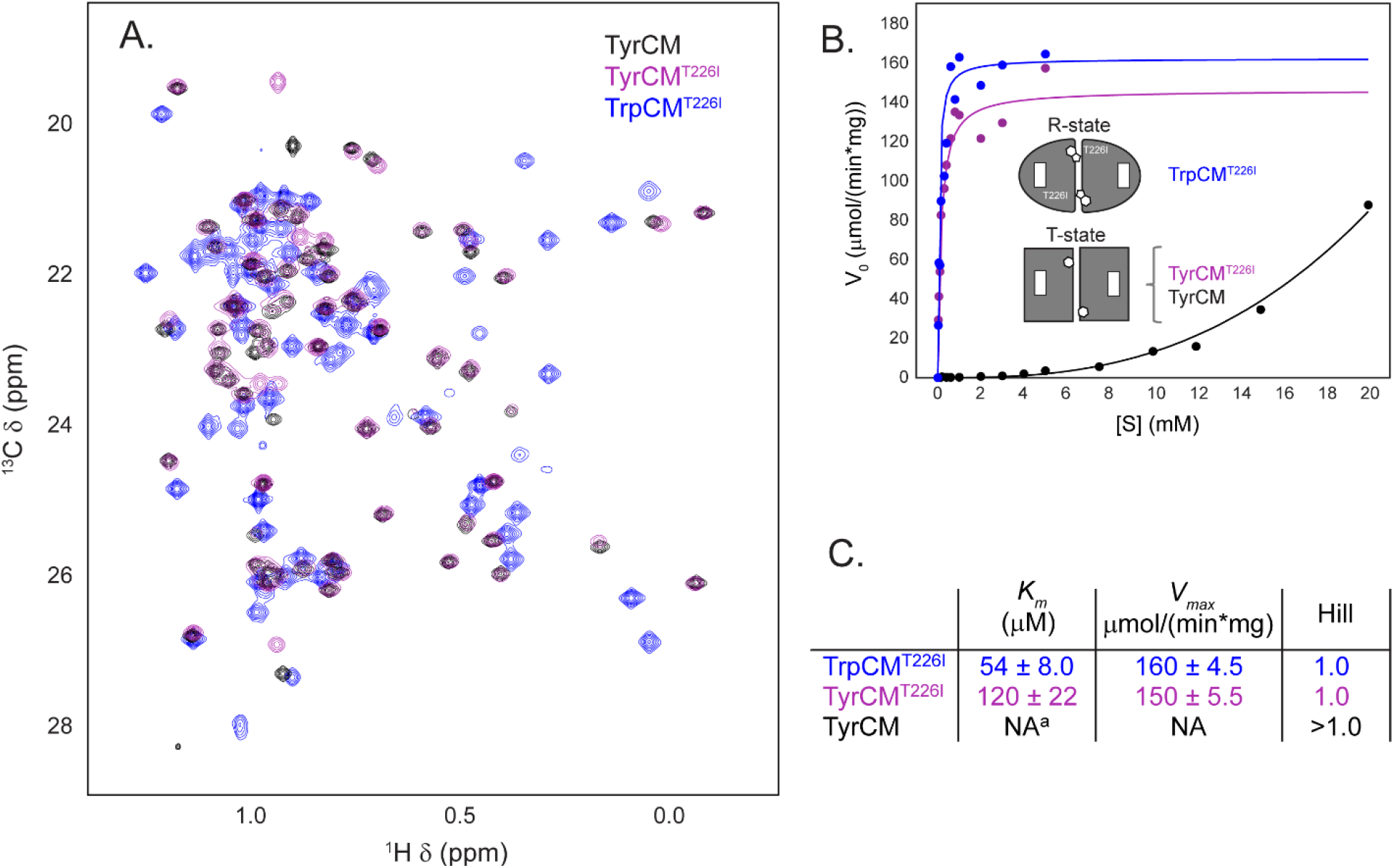
T conformation; maximum activity. (A) HMQC Leu/Val region of three CM forms, one in the R-state, and two in T. Wild type TyrCM is in the T state and fully inhibited while TyrCMT226I is also in the T-state, yet its activity (B & C) is virtually undisguisable from TrpCMT226I in the R conformation.

### Opposite trending with salt

TyrCM^T226I^ is one data point. Upon our previous characterization of the T-to-R switching of TrpCM in solution in which R is pre-sampled with an 8% population, it was evident that the dynamics of TrpCM we observed at 150 mM NaCl(23) differed from those observed at 0 mM NaCl(28). We confirmed this by measuring the dynamics of TrpCM at 0 M NaCl by methyl ^1^H CPMG (Figure 2A) and saw a sharp decrease in the population of the R conformation from 7.7 to 3.1% (Figure 2B). From the CS-based view, one would expect lowered activity at 0 mM salt relative to 150 mM in which the R conformation is favored. However, the opposite occurs: binding is *tighter* at 0 mM NaCl (Figure 2C). These data are also inconsistent with a mean population of high and low affinity states mechanism for substrate affinity.

**Figure 2.**
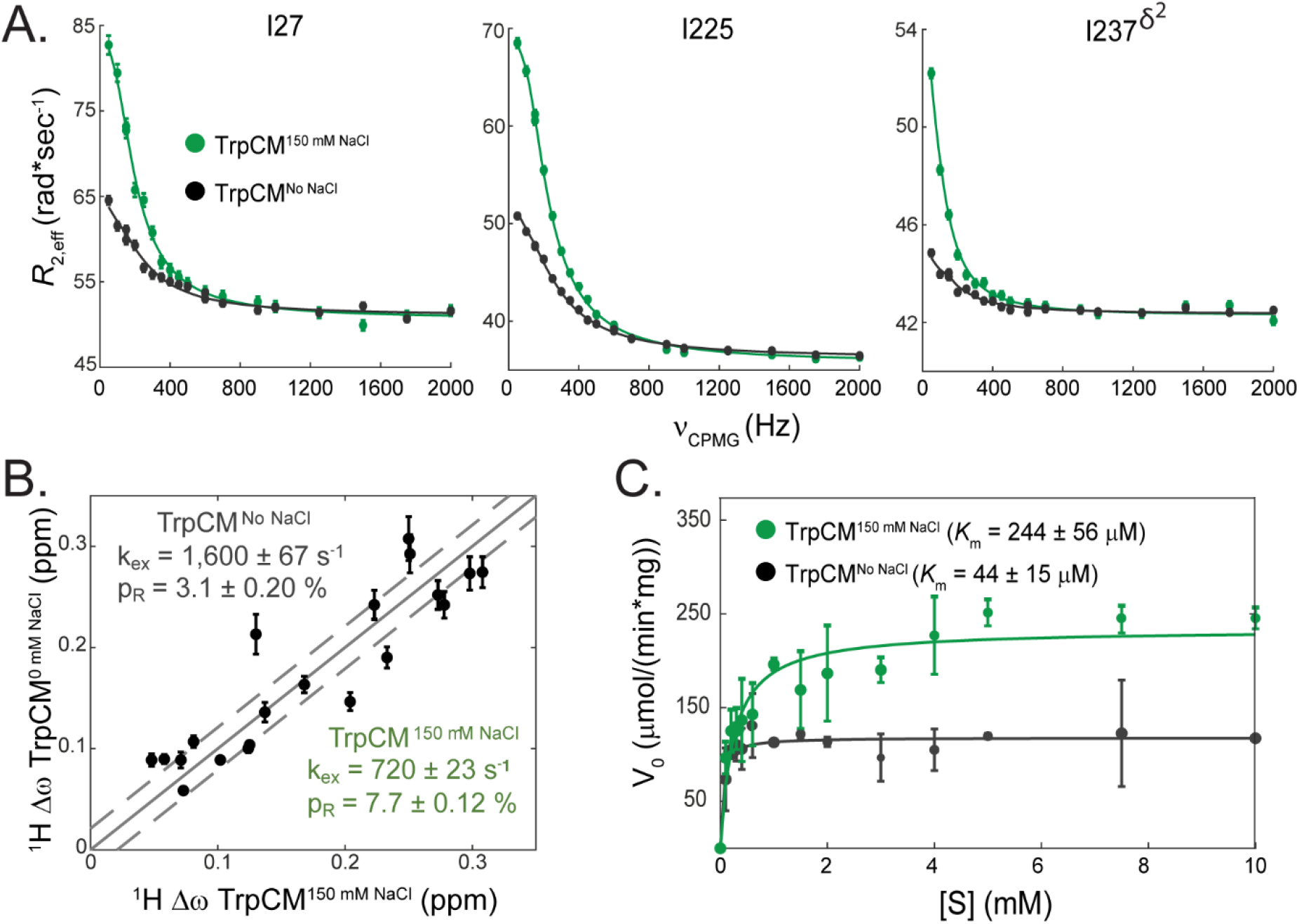
Reducing the population of R yields tighter substrate binding. (A & B) 1H ILV methyl CPMG shows salt promotes R-state occupancy. The population of R is decreased from 7.7 to 3.1% upon going from 0.15 to 0 M NaCl. (C) Reduction of pR is accompanied by a ∼5.5-fold increase in substrate affinity.

### R conformation, low activity

We recently showed that the D215A mutation in loop 11-12 of CM results in a substantially less-active enzyme under our standard condition of 150 mM NaCl, pH 7.5(29). In the context of bound activator (TrpCM), the double mutant D215A/T226I is observed to be primarily in the R conformation in the absence of active site ligand, as evidenced by methyl ^1^H-^13^C HMQC spectra. Most methyl peaks appear close to R-state peaks as defined by TrpCM^T226I^ (Figure 3A); however, many peaks diagnostic of T to R dynamics, such as I27 ^δ1^ and I225^δ1^ have disappeared, potentially due to an equilibrium with local conformations similar to the T-state. Even though the conformation of TrpCM^D215A,T226I^ is overwhelmingly R, the activity profile is low (Figure 3B). Thus, this form of CM provides the opposite situation in conflict with a CS mechanism to that shown above. These two CM forms together show contradiction with the T to R paradigm on both extremes of the conformation/activity scale.

**Figure 3.**
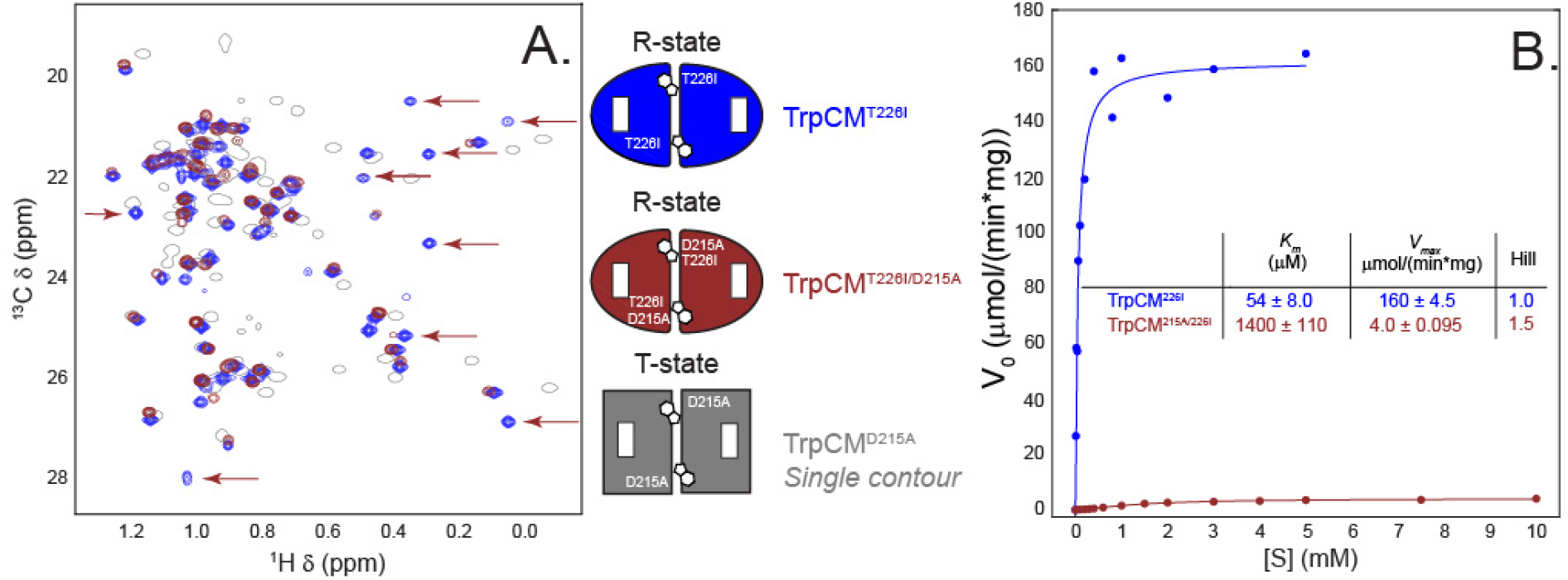
R-state conformation, weak binding. (A)TrpCMT226I and TrpCMD215A/T226I are in the R state as compared to TrpCMD215 which is in T. Note the missing peaks in the TrpCMD215A/T226I spectrum (arrows); these are the resonances with the largest Δω in the T vs. R forms, indicating that while the major state is clearly R, there is likely a sub-population of T (see text). (B) The R-conformation of TrpCMD215A/T226I does not lead to tight substrate binding.

### Poor correlation with a parameterized two-state T vs. R model

For an enzyme abiding by the two state CS-like paradigm, a correlation is expected between affinity (*K*_*app*_ or 1/*K*_*m*_) and the population of the high-affinity conformation (R) according to the normalized first order term of a binding polynomial giving the mean binding affinity for two interconverting species(30):

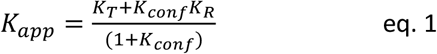

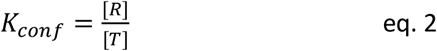

and *K*_*T*_ and *K*_*R*_ are the *K*_*m*_ values for the pure T and R states, respectively. (We note that in CM, *k*_cat_ is similar for apo, Trp, and TyrCM making *K*_m_ a good proxy for *K*_app_ in this system. Further chose the simplified, monomer type binding polynomial because activity profiles are hyperbolic—sites do not interact—when CM is bound to Trp(23)). We set out to compare our observations on CM within this framework. The affinity of the pure R-state, *K*_*R*_, was obtained from TrpCM^T226I^, whose spectrum is clearly R (Figure 1A), and for which relaxation dispersion shows little-to-no *R*_*ex*_ in resonances indicative of the T to R switch (Figure S4). The substrate affinity of the T-state is less straightforward because inhibited states tend to have positive cooperativity(24). In such cases the sigmoidal character and Hill fitting model dictate *K*_*m*_ is convoluted by two binding events: binding to the symmetrical inhibited T-state and binding to the activated single substrate bound protomer. To isolate the *K*_*T*_ for the first binding event, we created a mixed labeled dimer(31) with only a single catalytically competent subunit and measured its *K*_*m*_ (Supporting Methods and Figure S5), which is indeed weaker than any composite *K*_*m*_ observed and reported by us or in the literature, which was expected for a positively cooperative enzyme.

Simulated binding behavior for a two-state system, anchored by our measured pure CM *K*_*T*_ and *K*_*R*_ affinities is plotted in Figure 4A. The *K*_m_-*K*_conf_ pairs for the CM examples presented in this work all deviate significantly from the model, by nearly two orders of magnitude in some cases. The point for TrpCM^D215A/T226I^ requires further explanation: 1) While the spectrum shows the major state is R (Figure 3), many peaks that would report on T to R dynamics (arrows Figure 3) are broadened away, consistent with some population of T. To determine the relative populations, we acquired ^1^H CPMG data for this enzyme, and, although the rate of exchange from a global fit of a T to R sub-group was well determined (1100 ± 200 s^-1^), the data were too noisy to obtain either precise Δω values or the minor state population. We therefore made a conservative estimate of 85% R because some peaks in this group that are weak in TrpCM (8% minor state population) are fully broadened away in TrpCM^D215A/T226I^, consistent with greater than 8% of the excited state. 2) This is the only form in the plot that did not have a hyperbolic Michaelis-Menten curve. Therefor its x-coordinate in Figure 4A is based on a fitted composite *K*_*m*_ and Hill coefficient of 1.5 (Figure 3B). However, Hill>1 (positive cooperativity) means the fitted *K*_*m*,*obs*_ is *lower* (stronger binding) than the actual *K*_*m*_ for the first substrate binding event, and the data point would only be further below the conformational selection model line were we able to measure it directly (red point, Figure 4A). We note the binding affinities of TSI to apoCM vs TrpCM are dramatically different, in accordance with similar variance in estimated *K*_m_ values from activity assays (Figure S6), suggesting that *K*_m_ is a good proxy for *K*_D_. Taken together, these data are inconsistent with a two state model for the allosteric regulation of substrate binding in CM.

**Figure 4.**
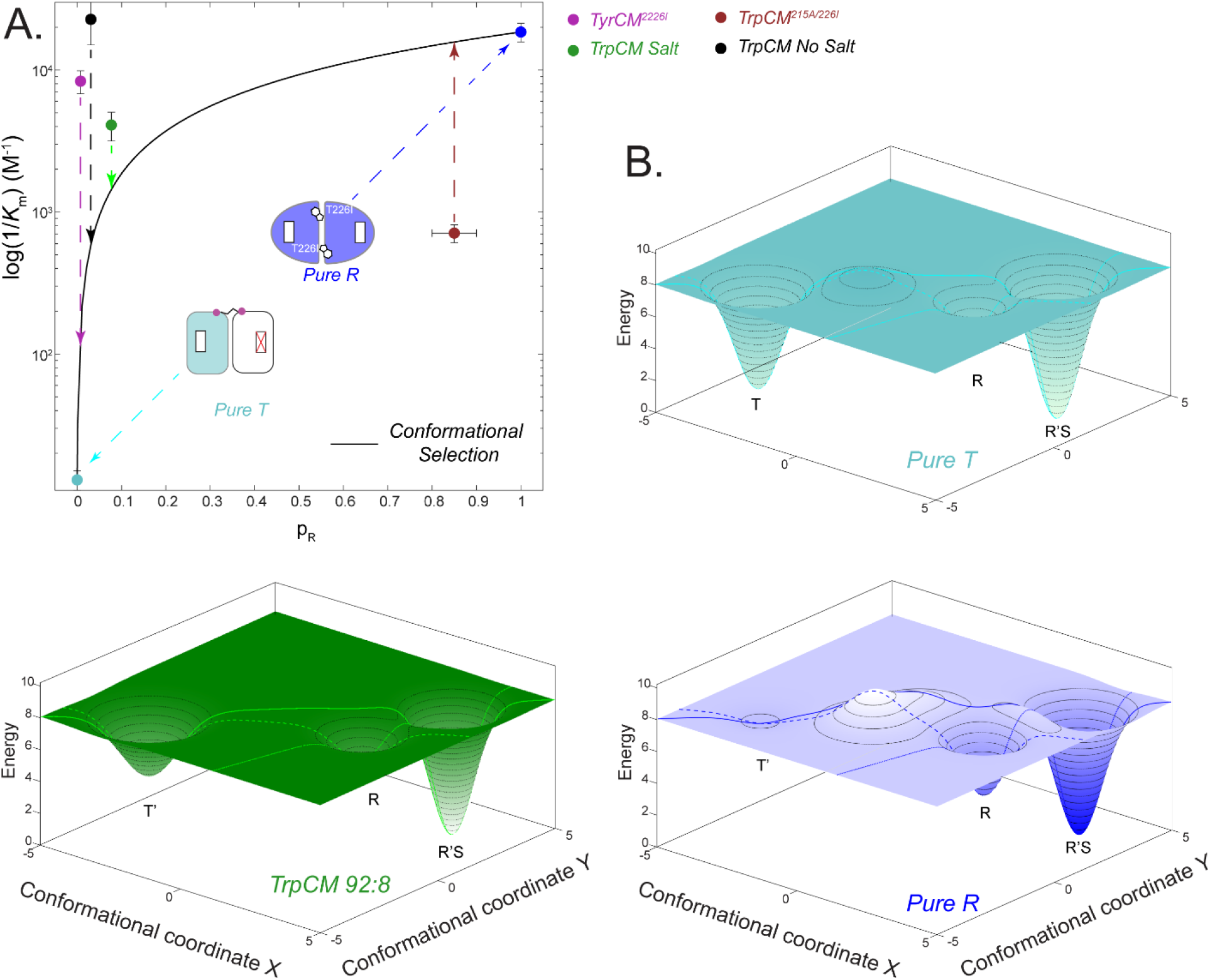
Range of K_m_ and R-state populations are incompatible with a conformational selection model. (A) K_m_ values for a series of CM states deviate from those predicted by CS by 1-2 orders of magnitude. (B) Energy diagrams for three states shown in panel A present a model for how R-state occupancy could be incidental. Each has three wells, the substrate-free T, the substrate free R, and the R’S (R’ is the “SuperR” conformer(42)). In the pure T state (cyan), the T basin is deep, and the R state is significantly disfavored. The TrpCM T state basin has higher energy, is wider, and has different coordinates than the pure T-state, so we denote it T’. This coupled with the lower energy of the R-state allows incidental escape from the basin and hence 8% pR. However, the increase in binding affinity is dominated by the large population and higher energy of the T’-state ensemble (green). The pure R TrpCMT226I enzyme exists entirely in the R conformer, but the change in energy between R and R’S is similar to that of T’ and R’S for TrpCM (blue).

## Discussion

The ability to clearly observe a ligand-bound-like protein conformation that appears spontaneously and transiently from an inactive conformation should be viewed as a technological achievement and a marker for how far structural biology has advanced. We previously observed this dynamic sampling in chorismate mutase bound to allosteric effector tryptophan (TrpCM) as a millisecond-timescale, two-state switching between T and R conformations, although the switch was not perfect and excluded some residues near the active site and within the effector binding domain(23).

Here, the larger body of evidence from multiple forms of CM indicates that this pre-formation of the active conformation (R) is not, on its own, functionally advantageous. In this case, and arguably in others, such motions appear to be incidental with respect to binding or other function. We are aware of at least five other reported instances of this. The first is catabolite activator protein (CAP), where multiple mutants occupied a high-affinity DNA binding conformation, in the absence of DNA, to different extents. However, the population of high-affinity states did not correlate with DNA binding affinity. Instead, the meaningful correlation with binding affinity was found to be with conformational entropy(13). The second is with the human form of thymidylate synthase, where switching from an active to an inactive form was detected(32) but could not provide a basis for CS-derived model of cooperative substrate binding(32, 33). Pyruvate kinase, another classically allosteric multimeric enzyme, initially believed to adhere to the TR framework(34), undergoes the R to T structural transition upon binding amino acids that fails to inhibit the enzyme(17). And lastly, aspartate transcarbamoylase, an enzyme that was initially thought to adhere to the MWC model(35), was found to be regulated by ‘breathing’ motions sampling a spectrum of conformational states between T and R(36). These examples are contrasted by studies where conformational selection is operative in binding or catalytic function(6, 37-39) or where the binding mechanism is mixed between CS and IF depending on the ligand concentration(20). It is notable that the most successful design of allosteric proteins took inspiration from the MWC formalism in the approach and the ensuing multimeric designs exemplified the key structural and functional features(40) envisioned 60+ years ago. Therefore, the main point is not to discount the power of the simple models, but rather to provide a vivid example that observation of a pre-sampled active-like conformation should not be interpreted as proof of a CS/MWC mechanism without further evidence. We note that the pioneering theoretical(18, 19) and experimental(20) work on flux based hybrid models was based on “simple” monomeric proteins. CM is an excellent test case to examine these ideas on an allosteric dimeric protein. However, fitting the additional myriad of kinetic rates to determine flux through the various pathways requires significant further method development (Figure S7).

Why would a protein execute a change to a distinct conformation if it is not going to take advantage of what is a sophisticated feat? We propose that incidental switching results from a protein’s evolution to enable the second conformation, which in the case of CM is the substrate/transition state-bound pose. A functioning protein needs to switch back and forth between the two structures so sequences are needed that can robustly reside in either and access them readily. This may be particularly true in allosteric oligomers such as CM where conformational switches might contribute to subunit cooperativity in addition to catalytic function. Put another way, the deep minima on the energy landscape could result in difficulty avoiding those passages to the active (or R) state that would be ingrained in the free energy surface. Hence, relatively small perturbations – including mutations – of any sort might modulate the energy landscape, to facilitate incidental population of some state (Figure 4B). Such temporary relocation into that conformation may be inconsequential and could allow other mechanisms of function to operate without penalty.

For CM, the question remains: what is the operative mechanism if not CS? Consider TyrCM^T226I^, which is in the T conformation yet is fully active just like TrpCM^T226I^ in the R-state. The location of T226 at the base of loop 11-12 might provide a clue to solve this paradox and hint at a *local* activation mechanism: Namely, there are different sub-ensembles of the highly flexible loop 11-12 in the active vs inactive state that either drive or inhibit substrate binding(29, 41) (Figure S1B). The data presented here suggest loop 11-12 dynamics are decoupled from switching between T and R. In other words, the ‘primed’ loop ensemble can exist in either the T’ (as in TrpCM or TyrCM^T226^) or R (as in TrpCM^T226I^) contexts. The question then becomes whether loop differences exist in the substrate-free or bound ensembles. Available evidence supports the former based on the following: 1) Loop 11-12 poses are different in substrate free crystal structures depending on the form but are always in the “super-R” pose whether Trp or Tyr are bound(42). 2) In accordance with (1), NMR spectra of effector-free, TyrCM, and TrpCM bound to TSI have nearly identical NMR spectra. 3) V_max_ remains similar in the different forms so the activation barrier is largely unaffected. Thus, any change in the free energy level of the Michaelis complex would require a similar shift in the transition state, which is possible but seems less likely than the alternative: that the primary change in free energy occurs in the substrate-free enzyme as schematized in Figure 4B. Nevertheless, speculations are separate from the main conclusion here, namely, that observation of an activated conformation of a protein in the absence of its binding molecule does not automatically implicate CS as the mechanism of action. And finer details of rugged ensembles will need to be understood to explain and design allostery in ways that go beyond two state switches.

## Supporting information

Supplementary material:Detailed materials and methods and supplemental figures 1-5 as referenced in the main text.

## Supplementary Material Description

Detailed materials and methods and supplemental figures 1-7 as referenced in the main text.

## Acknowledgments

This work was supported by NIGMS awards to Andrew L. Lee (GM127698 and GM144348). We would like to acknowledge Dr. Stuart Parnhum at the UNC-Chapel Hill Biomolecular NMR laboratory, which receives funding from the National Cancer Institute of the National Institutes of Health (P30CA016086). This study made use of NMRbox: National Center for Biomolecular NMR Data Processing and Analysis, a Biomedical Technology Research Resource (BTRR), which is supported by NIH grant P41GM111135 (NIGMS).

