## Supplementary material:Detailed materials and methods and supplemental figures 1-5 as referenced in the main text. for "Incidental conformational switching in an allosteric enzyme"

### Supporting Information

#### Supporting Methods

*<u>Protein expression and purification.</u>* U-[$^2$H,$^{15}$N], Ile-δ1-[$^{13}CH_3$], Leu-δ-[$^{13}CH_3$], Val-γ-[$^{13}CH_3$] CM was expressed using M9+ media(1) and purified as described(2).

<u>Mixed labeled dimer preparation.</u> S97C/C39S U-[$^2$H,$^{15}$N], Ile-δ1-[$^{13}CH_3$], Leu-δ-[$^{13}CH_3$], Val-γ-[$^{13}CH_3$] CM was reacted with 20-fold molar excess PEG3 Bismaleimide (BroadPharm) for 15 minutes on ice in NMR buffer (see below) without reducing agent. Unreacted linker was removed with a G25 chromatography step, 110% unlabeled K137C/R16L/N194A CM was then added and allowed to conjugate on ice for one hour. The R16L/N194A mutations render that subunit catalytically dead. The resulting MLD is covalently linked between the 97 and 137 positions. The slight excess of triple mutant enzyme ensures that the isotopically labeled and active subunit are "soaked up" and any unlinked triple mutant enzyme is quiet in both kinetic and NMR experiments. Purity was confirmed by SDS PAGE and HMQC.

<u>Activity Assays.</u> The continuous UV-based activity assay was performed as described(3), with the modification that a BioTek Epoch 2 plate reader was used to streamline and accelerate data acquisition. Data were acquired in NMR buffer, pH 6.5, 21 °C.

*<u>NMR Spectroscopy.</u>* HMQC spectra were acquired using an in-house coded pulse program on 850 MHz Bruker Avance III with TCI H-C/N-D 5 mm cryoprobes controlled by Topspin 3.5 pl 7. $^1$H CPMG experiments were acquired at 850 MHz and 600 MHz (TrpCM+salt and TrpCM$^{T226I}$), 850 MHz and 700 MHz (TyrCM$^{T226I}$), and 850 MHz only (TrpCM$^{D215A/T226I}$ and TrpCM no salt) as described(2, 4). Data were fit using Chemex (https://github.com/gbouvignies/ChemEx) installed on NMRBox(5). $\Delta\omega$ data were reported as absolute values rather than sign discriminated. All NMR experiments were acquired at 15 °C, 25 mM sodium phosphate buffer, 1 mM EDTA, 1 mM DTT, pD 6.1. Salt was omitted in one case to modulate the population of R in TrpCM (see main text).

<u>Isothermal Titration Calorimetry.</u> ITC experiments were carried out on a Malvern PEAQ-ITC calorimeter at 15 °C in NMR buffer with TCEP replacing DTT. The cell contained either effector free CM or CM in 5 mM L-Trp. The syringe was loaded with either TSI(6) or TSI in 5 mM Trp. Thermograms were integrated with NITPIC(7).

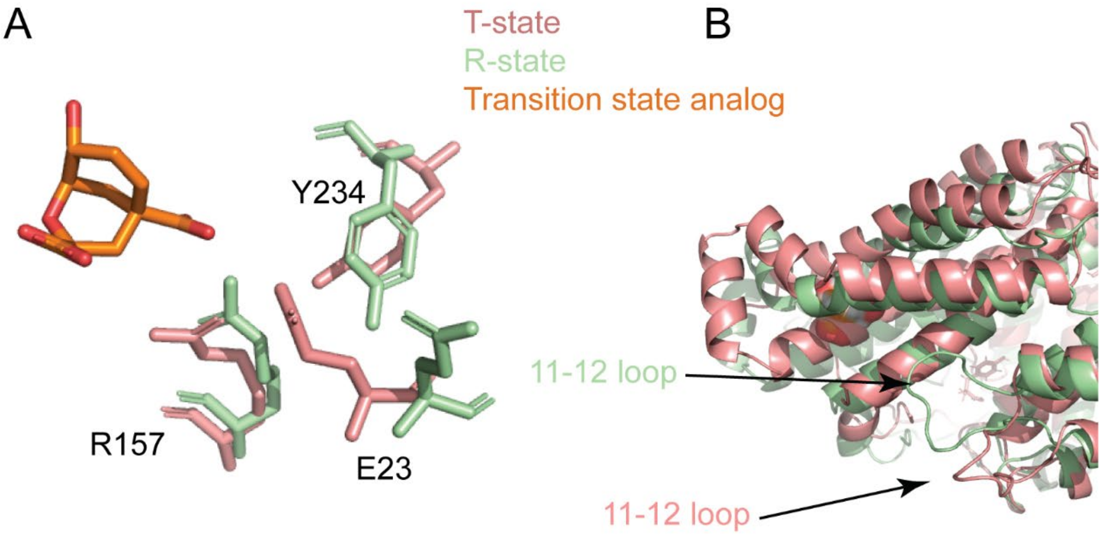


***Figure S1.*** A) The R-state x-ray model is expected to bind negatively charged substrate better than the T-state by virtue of relief of electrostatic (E23 moves away from binding pocket and R157 towards) and steric (Y234 rearrangement) repulsion. B) MD simulations (snapshots shown) and paramagnetic resonance experiments(8) show that loop 11-12 is also biased towards different poses in different effector-bound states. We hypothesize herein that this loop might adopt the activating pose in the absence of the rest of the T to R conformational changes.

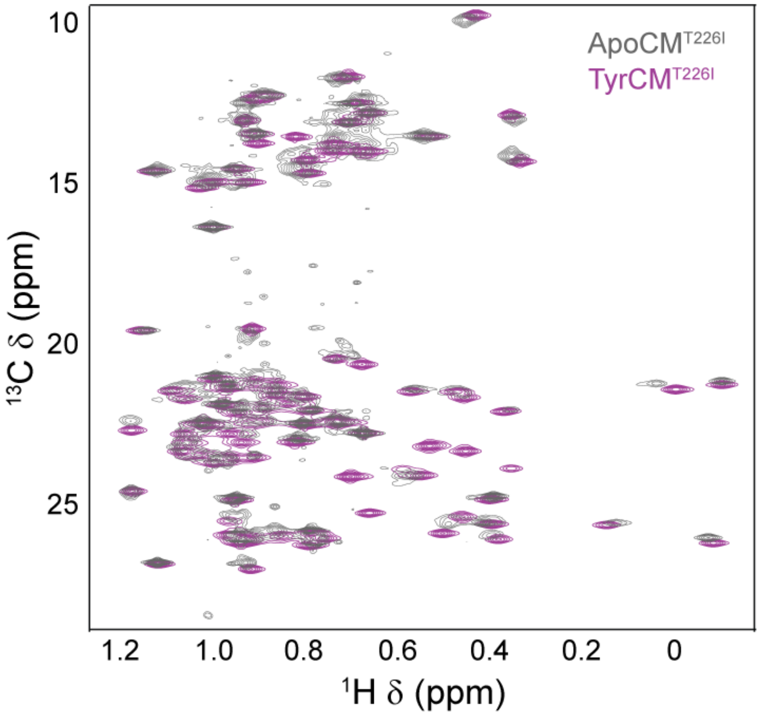


***Figure S2.*** Tyrosine binds to $CM^{T22I}$ which remains in the T-state. These data are required because it was previously reported by less reliable equilibrium dialysis experiments that Tyrosine fails to bind this mutant(9).

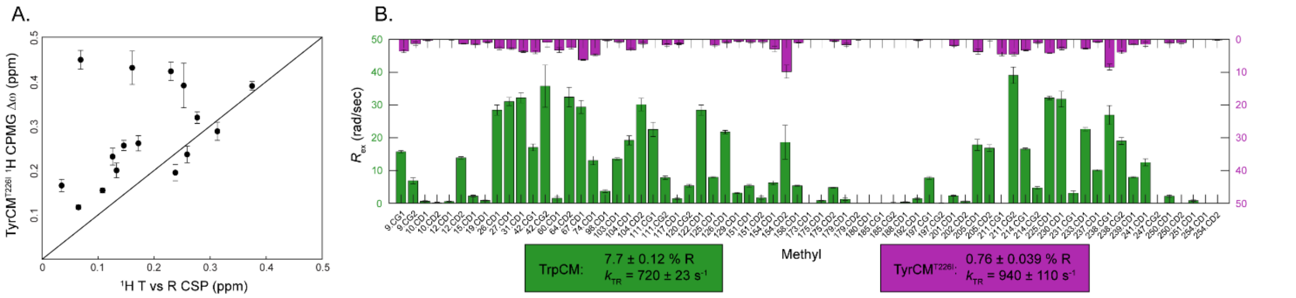


***Figure S3.*** TyrCM$^{T226I}$ samples the R-state, but the population is 10-fold lower than wild-type TrpCM. (A) $\Delta\omega$ values from a global fit of TyrCM$^{T226I}$ $^{1}$H CPMG curves correlate well with the difference in chemical shifts between the T and R states indicating that it samples the R-state in the absence of substrate. (B) Attenuated $^{1}$H CPMG $R_{ex}$ in TyrCM$^{T226I}$ vs. wild-type TrpCM yield fitted R-state population that is less than 1%. We show in the main text Figure 4 that TyrCM$^{T226I}$ actually binds substrate *tighter* than TrpCM despite the lower R-state population.

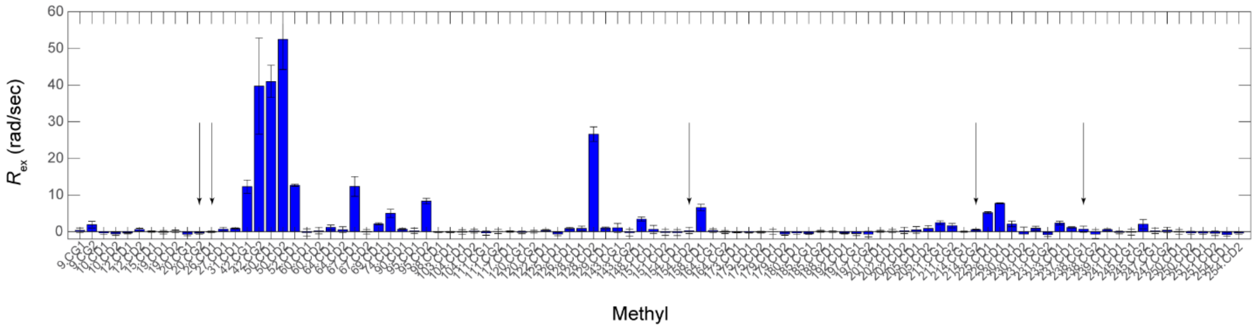


***Figure S4.*** TrpCM$^{T226I}$ is in the R state(10) and there is no evidence of switching to T. TrpCM$^{T226I}$ exhibits chemical exchange in the effector binding domain (defined as residues 42-104), and near loop 11-12, but none of the resonances most diagnostic of the T→R switch (highlighted with arrows) exhibited $R_{ex}$. As with other chemical exchange data described in this work, these values are from an ILV methyl $^{1}$H CPMG experiment.

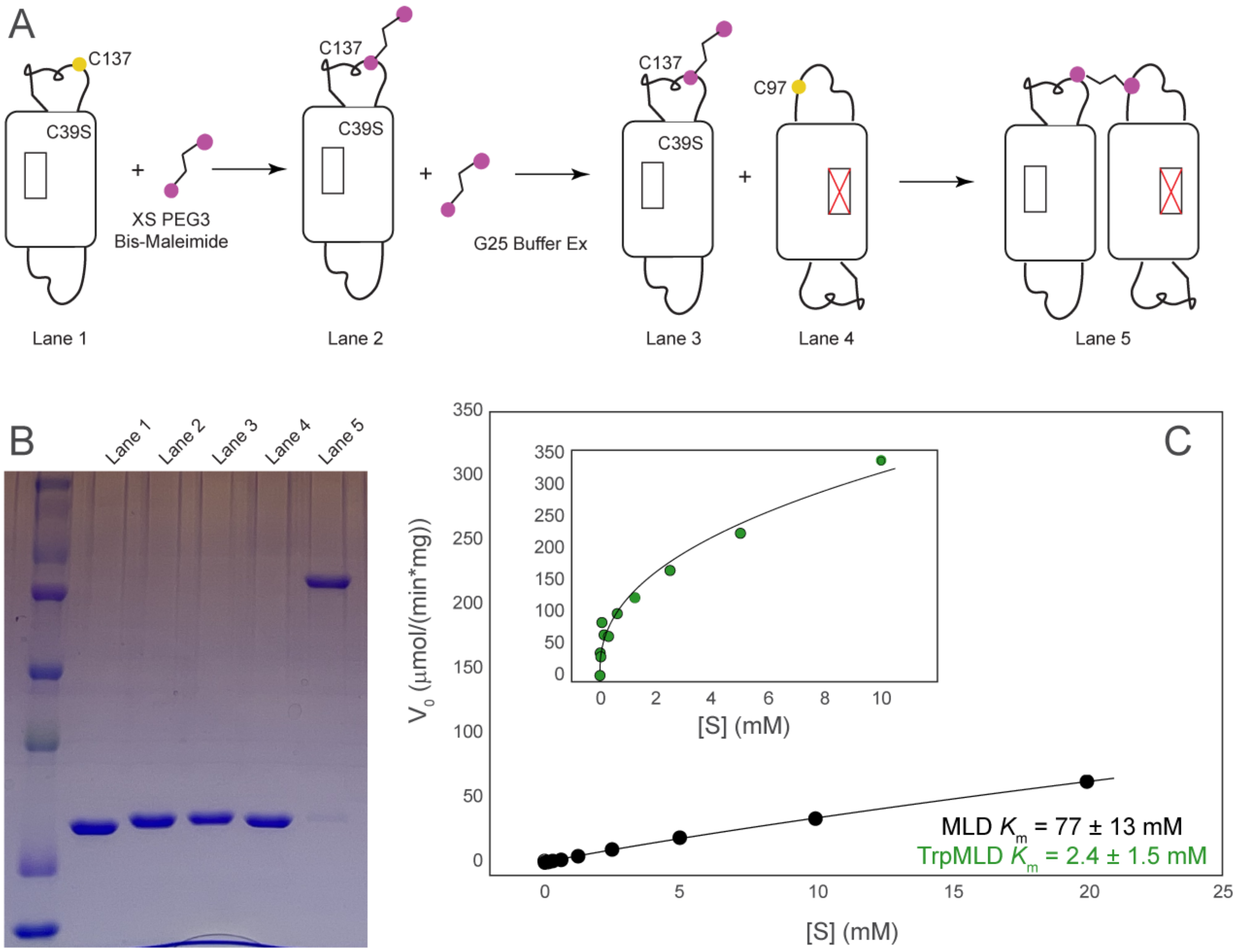


***Figure S5.*** (A) Mixed labeled dimer (MLD) strategy to extract the $K_m$ for the first T state substrate binding event. Inhibited forms of CM have positive cooperativity, therefore $K_m$ values from Hill-type models are convoluted by two binding events. To isolate binding to the first pure inhibited state we constructed a mixed labeled dimer with only one competent active site. The second active site was rendered inactive by a dual R16L/N194A mutation (denoted by X in panel A). Two different benign single cysteine mutants were engineered into a loop. One of the variants was then tagged with excess PEG3 bis-maleimide, the excess tag was removed, and the second mutant was spiked into the reaction with a stoichiometry to 'soak up' all the functional active site. The strategy was verified by SDS-PAGE (B) and NMR (one of the protomers was isotopically labeled and the other was not-data not shown because SDS-PAGE is unambiguous). (C) The $K_m$ of the effector free MLD was measured as a proxy for the isolated inhibited enzyme, as well as for the Trp-bound MLD to confirm it retained a wild-type-like effector profile, which it does.

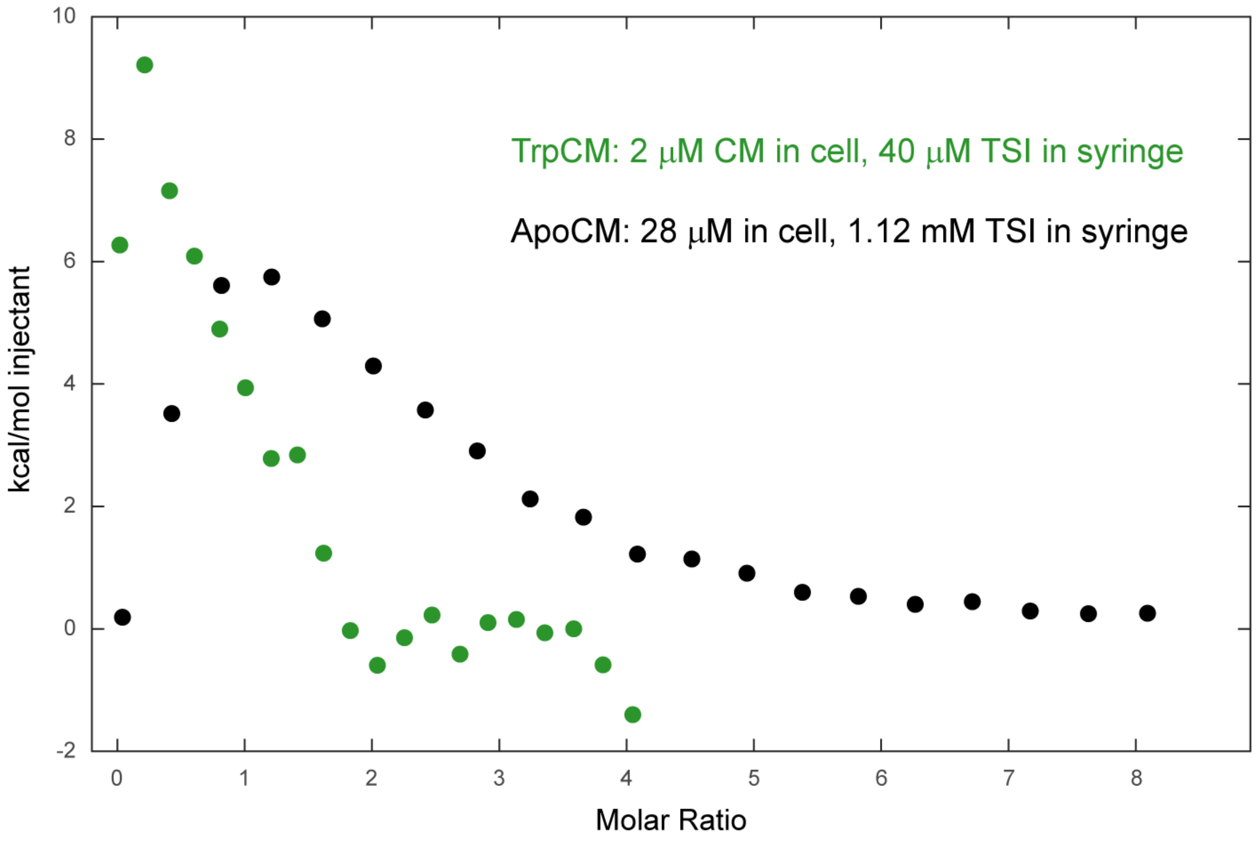


***Figure S6.*** Transition state inhibitor(6) (TSI) binding is also tighter to TrpCM than the apo enzyme. Given our(11, 12) and others reported difficulty in fitting multi-site binding with single concentration ITC experiments and the noise associated with low concentration/low heat in TrpCM experiments, for the purpose of this work, we rely on qualitative data. TrpCM saturates with TSI at a lower molar ratio despite being present at a 14-fold lower concentration in the cell and a 28-fold lower concentration in the syringe than the effector free enzyme setup. This supports two assertions: 1) $K_m$ is a good proxy for $K_D$, and 2) given similar $V_{max}$ values for the illustrative examples here, the differences in binding affinity are likely due to differences in the substrate-free enzyme ensembles.

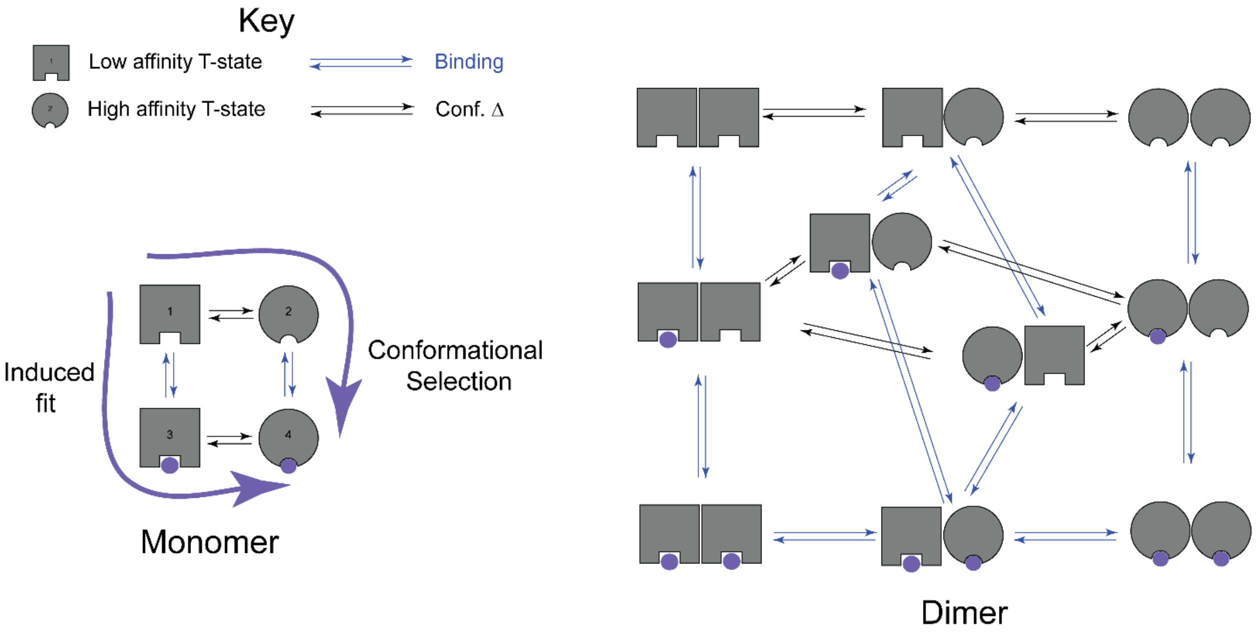


***Figure S7.*** Distinguishing between CS and IF flux through a dimeric enzyme is more than twice as complex as it is for a monomer. The schematic used to determine ligand flux through galectin-3 using state of the art approaches is shown on the left (adapted from (13)). A total of 7 kinetic rates were fit along with the assumption of diffusion limited on rates to both the high and low affinity states. The multiparametric fit was made possible by predetermined chemical shift changes for the binding and conformational change events. On the right, we show the complexity of the model balloons from 4 states to 10 states, and the number of rates proliferates even further. In addition, the architecture of the dimer is such that there are not methyl chemical shift probes at the dimer interface(2) to detect mixed states. We are in the process of using mixed labeled dimers to untangle this question (see figure S4 above).
